# IQCELL: A platform for predicting the effect of gene perturbations on developmental trajectories using single-cell RNA-seq data

**DOI:** 10.1101/2021.04.01.438014

**Authors:** Tiam Heydari, Matthew A. Langley, Cynthia Fisher, Daniel Aguilar-Hidalgo, Shreya Shukla, Ayako Yachie-Kinoshita, Michael Hughes, Kelly M. McNagny, Peter W. Zandstra

## Abstract

The increasing availability of single-cell RNA-sequencing (scRNA-seq) data from various developmental systems provides the opportunity to infer gene regulatory networks (GRNs) directly from data. Herein we describe IQCELL, a platform to infer, simulate, and study executable logical GRNs directly from scRNA-seq data. Such executable GRNs provide an opportunity to inform fundamental hypotheses in developmental programs and help accelerate the design of stem cell-based technologies. We first describe the architecture of IQCELL. Next, we apply IQCELL to a scRNA-seq dataset of early mouse T-cell development and show that it can infer *a priori* over 75% of causal gene interactions previously reported via decades of research. We will also show that dynamic simulations of the derived GRN qualitatively recapitulate the effects of the known gene perturbations on the T-cell developmental trajectory. IQCELL is applicable to many developmental systems and offers a versatile tool to infer, simulate, and study GRNs in biological systems. (https://gitlab.com/stemcellbioengineering/iqcell)

## INTRODUCTION

Stem cell fate decisions are made via dense arrays of interacting transcription factors (TFs) forming gene regulatory networks (GRNs) (Semrau & van Oudenaarden, 2015). Information gleaned from GRNs in stem cell differentiation can lead to more effective design-based cell cultures, applicable to cell therapies (Lipsitz et al., 2016; Prochazka et al., 2017). As a prominent example, the effect of transcription factors on GRNs has been widely utilized in the reprogramming of embryonic and adult somatic cell GRNs for the establishment of a pluripotent state via induction of driver TFs (Takahashi & Yamanaka, 2006). Stem cell reprogramming and differentiation can be modeled as executable and logical (Boolean) GRNs undergoing state transition (Sara-Jane Dunn et al., 2019; Peter et al., 2012; Yachie−Kinoshita et al., 2018). Executable GRNs provide information about both the topology and the regulatory rules of gene interactions that can be simulated as time-evolving (dynamical) systems. However, deriving informative, executable, and predictive GRNs for stem cell differentiation has proven to be a challenging task. Specifically, developing executable GRNs by piecing together evidence from gene perturbation experiments has shown to be an effective strategy (Peter et al., 2012) but is extremely time-consuming, labor intensive, and expensive. In a notable advancement, automated formal reasoning successfully identified a set of minimal GRNs underlying naive pluripotency in mice. Gene expression observations across multiple culture conditions were used to logically constrain possible GRN configurations, and the resulting set was able to accurately predict the outcome of 70% of new experiments(S.-J. Dunn et al., 2014; Yordanov et al., 2016). Yet, these methods are not based on high-throughput data.

More recently, the emergence of single-cell profiling technologies has provided an unprecedented archive of information regarding cells undergoing fate determination and maturation.. Deriving more accurate GRNs based on sc data is at the center of many recent efforts (Babtie et al., 2017; Fiers et al., 2018; Pratapa et al., 2020). Formal reasoning has been used to infer executable GRNs directly from high throughput single-cell quantitative PCR (sc qPCR) data (Hamey et al., 2017; Moignard et al., 2015). However, using single-cell RNA-sequencing (scRNA-seq) data has many advantages in terms of coverage, availability, flexibility in gene selection, and accuracy in clustering and pseudo-time inference compared to sc qPCR. These benefits exist alongside the disadvantage of dropout effects and low sensitivity in profiling TFs. Despite the availability of this data resource, this emerging field is still missing an integrated platform to infer, study, and simulate executable GRNs directly from scRNA-seq.

Herein we report an effective strategy implemented in a Python software package (IQCELL) for reconstructing GRNs directly from scRNA-seq data, called IQCELL. Our method includes steps for correcting dropout effects, selecting desired genes, building logical GRNs directly from pseudo-time with respect to interaction hierarchy and mutual information between gene pairs, and simulating developmental trajectories under normal and perturbed conditions. We demonstrate the utility of IQCELL by reconstructing a GRN for early mouse T-cell development, a well-characterized mammalian developmental system (Longabaugh et al., 2017), using published scRNA-seq data (Zhou et al., 2019). Our resulting GRN recovers over 75% of experimentally validated causal gene-gene interactions spanning years of research. Dynamic simulations of the inferred GRN resemble experimentally observed gene expression dynamics and capture the effects of knocking out or forcibly expressing various genes during early T-cell development. Our method is generally applicable to scRNA-seq data of differentiating cells and should serve as a useful resource for the community.

## RESULTS

### Integrative Method for Predicting the Qualitative Effect of Gene Perturbations on Developmental Trajectories of Cells (IQCELL)

IQCELL infers logical regulatory networks directly from existing information in the scRNA-seq data of cells during development and uses these regulatory networks to simulate and predict the behavior of developing cells under perturbed conditions (**Figure 1**). IQCELL works with quality controlled and pre-processed scRNA-seq gene expression data (Butler et al., 2018; Wolf et al., 2018). The second input of the IQCELL platform is the inferred pseudo-time ordering of cells based on scRNA-seq data (**Figure 1**). The temporal dynamics of genes helps with the inference of causal gene interactions. Pseudo-time ordering of cells using scRNA-seq data has been shown to be informative for capturing temporal and developmental dynamics (Haghverdi et al., 2016; Qiu et al., 2017).

**Figure 1.**
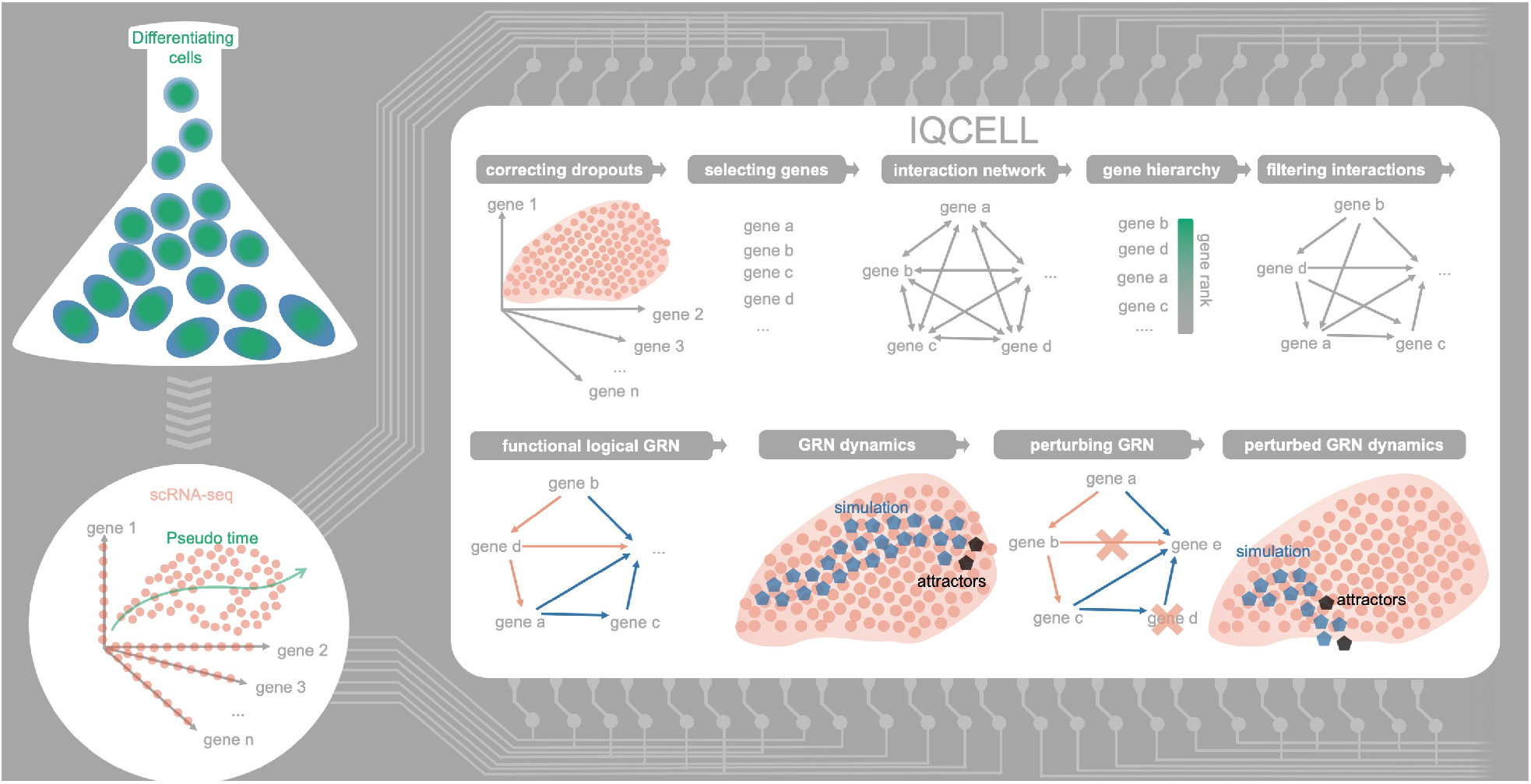
Overview of IQCELL. IQCELL infers logical GRNs directly from sc-RNA seq data and allows the simulation and analysis of *insilico* developmental trajectories in normal and perturbed conditions. The typical inputs of IQCELL are sc-RNA seq expression data along with the pseudo-time ordering of the cells. After correction of dropout effects and gene selection steps, gene-gene interactions are calculated and weighted based on mutual information. Binarized gene expression values are used to constrain possible gene-gene interactions and obtain a functional GRN for the data. IQCELL can be used to analyze the GRN and simulate possible developmental trajectories under normal and perturbed conditions.

Since gene dropout is common among many scRNA-seq datasets, particularly for transcription factors with low mRNAs copy numbers, IQCELL employs a recently developed graph-based algorithm (MAGIC) to recover gene expression (van Dijk et al., 2018) (**Figure S1A**). After selecting genes of interest based on literature curation (see **Figure S1B** for more details), we generate a set of possible interactions between genes. Information-based metrics such as mutual information are well-suited for quantifying relationships between genes (Song et al., 2012). IQCELL scores gene-gene interactions according to the mutual information between gene pairs (Krishnaswamy et al., 2014) and assigns a regulatory sign (activation or repression) to each interaction based on the significance and sign of their correlation (**Figure S1C**). These steps result in a dense weighted network of gene-gene interactions that needs to be filtered into a functional GRN. In a functional GRN, interactions are not necessarily biophysically direct but capture the consequence of regulatory relations.

To reduce the number of possible gene interactions, IQCELL forms a gene interaction hierarchy in which higher ranked genes influence lower ranked ones. To form this hierarchy, IQCELL binarizes the gene expression counts by clustering them into expressed and non-expressed states (Macqueen, 1967). The binarization process divides the pseudo-time axis into regions with compact and sparse expression densities for the genes, reflecting the pseudo-time domains where a gene is expressed at a higher or lower level (**Figure S1D**). Next, the platform identifies the transition points between expression regions for all genes and uses the order of transitions to form a gene interaction hierarchy, with highly ranked genes (with earlier transition points) having greater potential to influence those downstream. This acts as an additional filter on gene-gene interactions along with the mutual information (**Figure S1E**). The resulting directional network serves as the foundation for inferring executable GRNs.

To obtain an executable GRN model, IQCELL models interactions between genes as Boolean logic functions (Yachie−Kinoshita et al., 2018). IQCELL uses a satisfiability modulo theory engine (Z3) (**Figure S1F**), (de Moura & Bjørner, 2008) to identify logic functions that are compatible with the pseudo-time dynamics of binarized gene expression states.(Hamey et al., 2017). Finally, it selects the GRN with the highest average mutual information as the most probable constrained model. The result is a functional and executable GRN that optimally fits the input scRNA-seq data. IQCELL has built-in capabilities to simulate GRN dynamics via random asynchronous Boolean simulation (Yachie−Kinoshita et al., 2018) and compare the results with experimental data under normal and perturbed conditions. In summary, IQCELL processes scRNA-seq data inputs to infer an executable logical GRN that best fits the data.

### IQCELL sorts genes based on transition points and places them in a biologically relevant order

To assess the functional capabilities of IQCELL, we evaluated its performance using a well characterized mammalian developmental system, mouse early T-cell development (Hosokawa & Rothenberg, 2020; Yui & Rothenberg, 2014). The T-cell developmental program takes place in the thymus. It drives pre-thymic progenitors differentiated within the bone marrow toward T lineage commitment, and involves dense network of genes (Kueh & Rothenberg, 2012). Sustained exposure to Notch signaling drives early thymic progenitors (ETP) to the *CD4/8* double-negative 2A (DN2A) and DN2B stages, where upregulation of T-cell lineage-specific genes and progressive loss of potential for other blood cell fates occurs. Once committed to the T-cell fate, the double-negative 3 (DN3) T-cell progenitors begin recombining the β-chain of the pre-T-cell receptor (TCR). Cells are selected for functional β-chain rearrangements through pre-TCR signaling and proceed toward *CD4/8* double-positive state (DP) (**Figure 2A**). We used a publicly available scRNA-seq dataset (Zhou et al., 2019), where the authors used fluorescence-activated cell sorting to capture mouse thymocytes at ETP-DN2 and DN3 stages based on cell surface markers. After processing the data in a manner consistent with the original publication (**Figure S4** and **Table S2**), the gene expression profiles and the pseudo-time orders were used as inputs to IQCELL.

**Figure 2.**
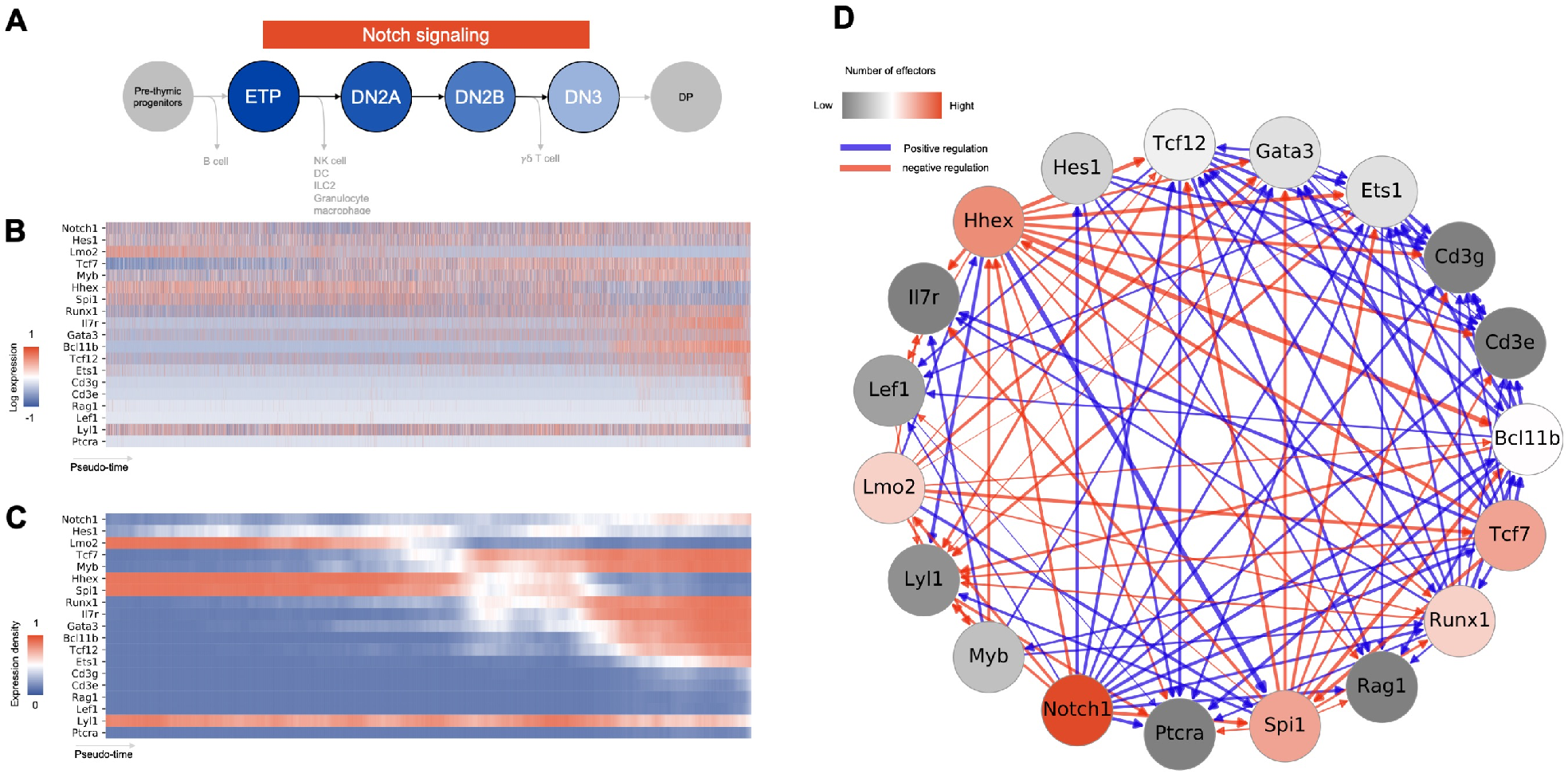
IQCELL initial processing of early T-cell development scRNA-seq data. (A) Summary of the scope of the scRNA-seq data used as an input to IQCELL (Zhou et al., 2019). ETPs originated from pre-thymic progenitors progress toward DN2A, DN2B (coincides with upregulation of Bcl11b and lineage commitment), DN3 stages and eventually lead to DP cells (not covered here). (B) Log transformed expression matrix for selected genes from scRNA-seq data along the pseudo-time axis. Gene expression is corrected for dropout effects using MAGIC (van Dijk et al., 2018).. Red indicates high expression, blue indicates low expression. (C) Smoothed binarized gene expression matrix (expression density). Gene expression values were binarized by clustering, averaged over a pseudo-time window, then sorted based on transition points from early to late. Red indicates high expression, blue indicates low expression. (D) The set of all possible gene-gene interactions, filtered by interaction hierarchy and mutual information (and signed by correlation. Positive and negative interactions are represented by blue and red edges, respectively. Edge width represents the relative amount of mutual information of the interaction. Nodes colored red have higher out-degrees.

After expression recovery, selecting genes of interest based on expression variation and biological relevance (**Table S1** and **Figure 2B**), and finding possible gene-gene interactions, IQCELL binarized gene expression values and calculated the expression density over pseudo-time (**Figure 2C**) (see STAR Methods). Sorting the genes based on their transition points placed *Notch1* and *Hes1* at the top of the gene interaction hierarchy (since their expression level stayed relatively high consistently) followed by *Lmo2, Tcf7, Myb*, and *Runx1* which agrees with their position in the regulatory hierarchy during T-cell lineage establishment (Yui & Rothenberg, 2014); whereas DN3 associated genes such as *Cd3e, Lef1*, and *Ptcra* (Masuda et al., 2007; Yui et al., 2010) appeared at the bottom of the hierarchy (**Figure 2C**).

Next, IQCELL used this order of genes (**Figure 2C**) as a hierarchical filter of possible interactions, with the genes at the top having the most regulatory potential in terms of number of genes they can regulate. The combination of the regulatory potential of individual genes, the mutual information between gene pairs, and interaction signs, led to a directed gene interaction network comprising the set of possible interactions (**Figure 2D**). This network then constitutes a foundation for further constraints and analyses at next steps.

### IQCELL is highly predictive for functionally regulatory interactions

Following our *in silico* analysis, we compared predictions from our initial inferred interaction network to validated regulatory interactions in mouse T-cell development. The initial interaction network (**Figure 2D**) was simplified with additional constraints. These constraints enforce the gene interactions to follow the expression patterns throughout the pseudo-time axis (**Figure 2B**) when executed as a logical network. This resulted in a set of possible update logical rules for each gene, selecting the most probable interactions as scored by mutual information leading to a provisional executable GRN for early T-cell development (**Figure 3A** and **Table S1**).

**Figure 3.**
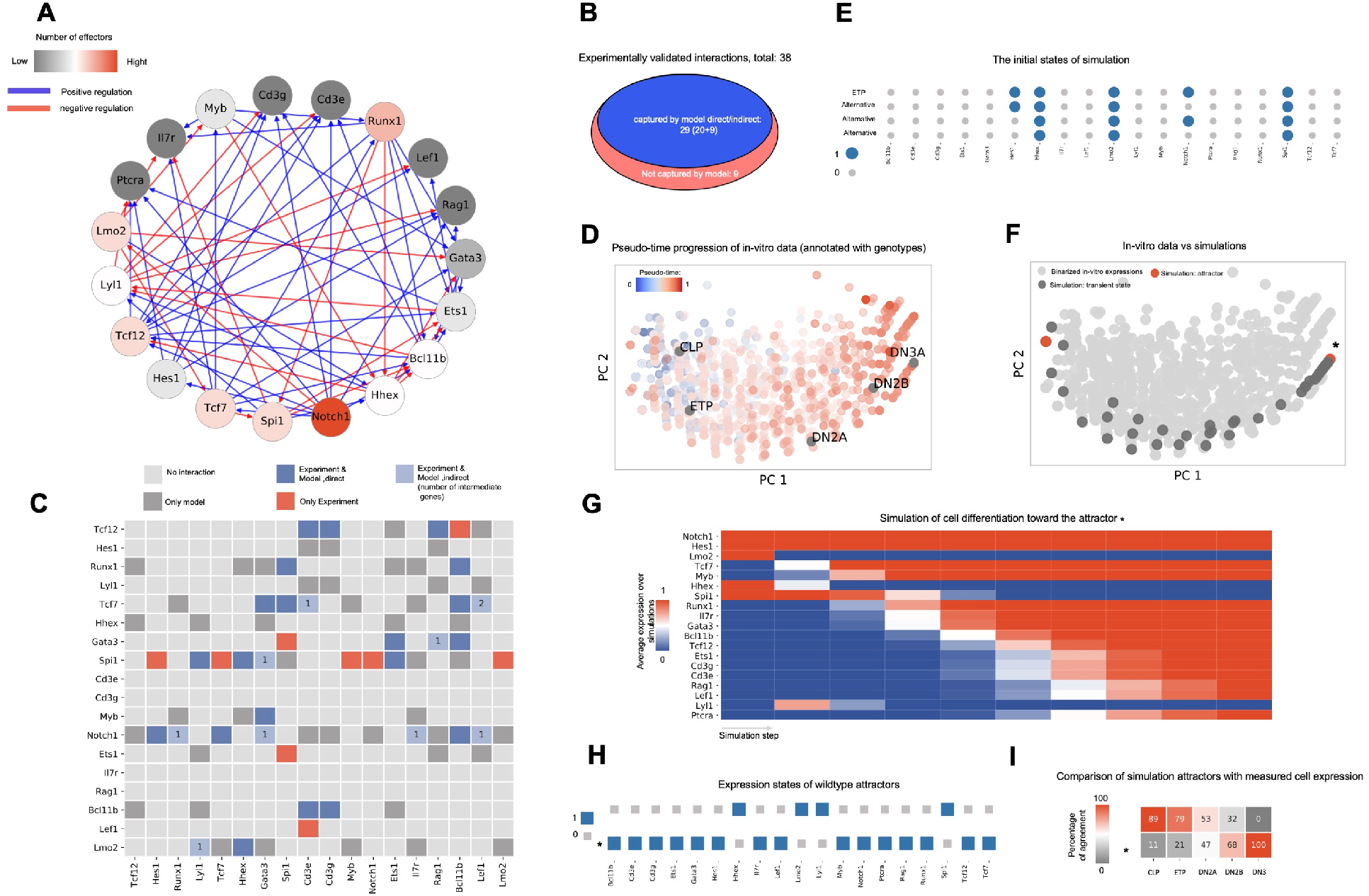
The provisional GRN for mouse early T-cell development inferred by IQCELL captures essential gene interactions and accurately simulates T-cell developmental trajectories. (A) The provisional GRN for early mouse T-cell development. The GRN is obtained by constraining the possible interactions to both follow the *in vitro* data progression when executed as a logical network and maximize mutual information between gene pairs. Positive and negative interactions are represented by blue and red edges, respectively. Nodes colored red have higher out-degrees. (B) Out of 38 experimentally reported gene interactions of early mouse T-cell development (Longabaugh et al., 2017), 29 of them are captured by the functional GRN model proposed by IQCELL. (C) Detailed representation of the proposed interactions by IQCELL and experimentally reported ones. Rows and columns represent regulators and effector genes, respectively. Blue indicates that the interaction is captured by the model directly (dark blue) or indirectly (light blue); in the latter case, the numbers indicate the number of intermediate genes. Dark gray indicates that the interaction is only proposed by IQCELL. The red color indicates the experimentally validated interaction is not present in the model. Light gray cells indicate no interaction. Genes downstream of *Spi1* comprise 50% of the experimentally-reported interactions not captured by IQCELL. (D) The PCA plot of the binarized scRNA-seq data color-coded with the pseudo-time values attributed to each cell. The binarization is performed by clustering the scRNA-seq expressions into expressed or not expressed levels. On top of that, the binarized expressions of CLP, ETP, DN2A, DN2B, and DN3A cells have been calculated from the immgen microarray data (Jojic et al., 2013) and overlaid on RNA-seq data. (E) The four initial states that have been used in simulations. Three variations of the state representing ETP are due to the noisy expressions of Notch1 and Hes1 genes in recovered sc-RNA seq data with early pseudo-time. Genes that are expressed (1) and not expressed (0) are represented with blue and grey circles, respectively. (F) The PCA plot of the simulated developmental trajectories are overlaid on the binarized scRNA-seq. The two detected attractors are colored red, and the attractor that matches the DN3A state is marked by star (*). (G) Average gene expression at each simulation step. All simulations started from the same initial condition (ETP) and move toward the same attractor (*). (H) Expression states of the GRN model steady state attractors. Genes that are expressed (1) and not expressed (0) are represented with blue and grey squares, respectively. (I) Percentage of similarity between the two attractors (vertical axis) and binarized microarray expression profiles of CLP, ETP, DN2A, DN2B, and DN3A cells (horizontal axis) (Jojic et al., 2013). The average agreement between two random states is 50%.

To benchmark this GRN, we compared our predicted directional interactions with a recent comprehensive GRN model of mouse T cell development based on experimentally validated gene interactions (Longabaugh et al., 2017). This network consists of 38 reported interactions between the genes of interest, of which 29 (over 75%) are de novo captured directly by our simulated functional regulatory network (**Figure 3B** and **Figure 3C**). For example, it is well known that *Bcl11b*, the gene that marks T lineage commitment, is activated by Notch signaling, *Gata3, Tcf7*, and *Runx1* (Kueh et al., 2016). Our model predicted four activators for *Bcl11b, Notch1* (as a Notch signaling mediator (Radtke et al., 1999) and target gene (Weerkamp et al., 2006) functionally represents the presence of Notch signaling), *Gata3, Tcf7*, and *Runx1*. The presence of *Runx1* in our model is a notable result. *Runx1* is present in developing cells, however it only gains access to the *Bcl11b* locus after chromatin restructuring during the DN2a stage (Ng et al., 2018). Notably, more than half of the interactions that were not captured by our model are related to the *Spi1* gene which is a T-cell lineage suppressor (Yui & Rothenberg, 2014).

To test the IQCELL performance with another source of data of T-cell development under different condition, we performed scRNA-seq analysis of *in-vitro* differentiation of fetal liver hematopoietic progenitor cells toward the T-cell lineage. Application of IQCELL to this second scRNA-seq data set provided further validation of its ability to predict gene-gene interactions (**Figure S3**).

One advantage of logical GRN models is that they can not only provide information about gene interactions, but can also be simulated to predict how the system evolves in time. To demonstrate this capability, we simulated our inferred logical GRN model and compared its output to experimental observations of mouse T cell development. The scRNA-seq expression data (Zhou et al., 2019) was binarized by grouping the gene expression count into on and off states. This data was then used in principle component analysis (**Figure 3D**) and the simulated trajectories overlaid on top of the binarized scRNA-seq gene expression data (**Figure 3F**). As the initial states of the simulations (representing the starting expression state of simulations), we used the binarized representation of cells at the beginning of the pseudo-time trajectory. These cells resemble the known expression state of ETP cells (Yui & Rothenberg, 2014). However, given the noisy expression of *Notch1* and *Hes1* at the earlier pseudo-time points (**Figure 2C**), we considered the expression states of these two genes to be random which results in four distinct initial states in total (**Figure 3E**). Two steady states have been obtained for the given initial cell states (**Figure 3F**), with one of them matching the DN3a cell profile (noted by star * in Fig 3). The simulated gene expression dynamics from ETP state towards this steady state shows a similar trajectory compared to the one observed from scRNA-seq data (**Figure 3G**, compare with **Figure 2C**). The other steady state shares similarities with common lymphoid progenitors (CLP) and ETP cells (**Figure 3H** and **I**). Overall, this analysis demonstrates that our GRN model is informative about both gene interactions and the behavior of genes at the system level. Such a model has the potential to predict the effect on gene perturbations at the system level as well.

### IQCELL predicts the effect of gene perturbations on developmental trajectories

Next, we tested the effect of gene perturbations on simulated developmental trajectories (**Figure 4A**). In particular, we tested the effect of gene perturbations known to result in halting or promoting T-cell development during the ETP-DN3 stages (reviewed in (Yui & Rothenberg, 2014)).

**Figure 4.**
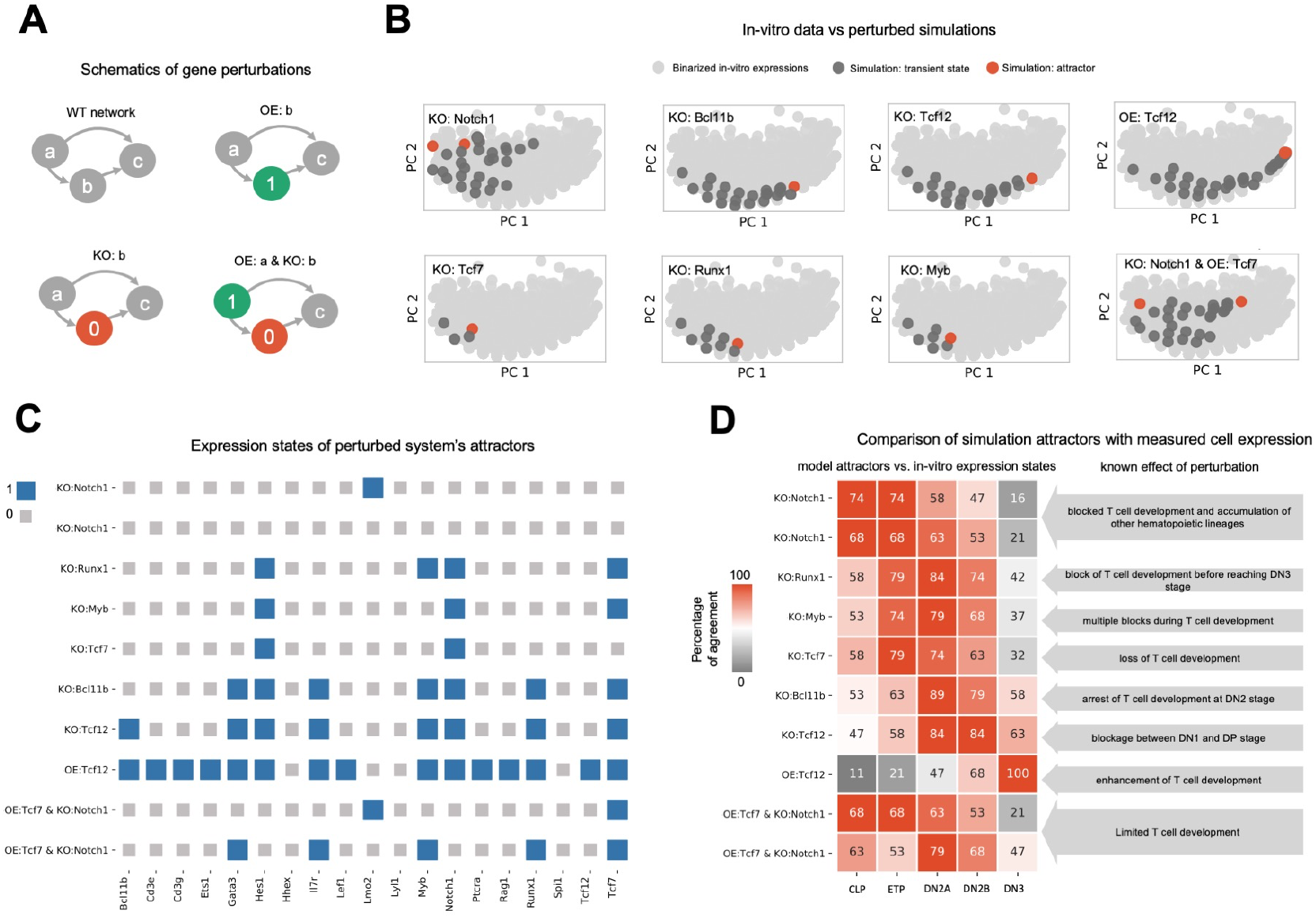
Testing the known effect of eight gene perturbations on in-silico developmental trajectories. (A) Schematic of performed gene perturbations. In overexpression (OE), the gene is always expressed (represented with 1) and in knockout(KO), the gene is always silent (represented with 0). (B) PCA plot of the simulated developmental trajectories under perturbed conditions are overlaid on the binarized scRNA-seq. The perturbations include knock-out of *Notch1*, knock-out of *Tcf7*, knock-out of *Bcl11b*, knock-out of *Runx1*, knock-out of *Tcf12*, knock-out of *Myb*, overexpression of *Tcf12* and the double perturbation, overexpression of *Tcf7* and knock-out of *Notch1* at the same time. (C) Expression states of the model attractors under perturbations. Genes that are expressed (1) and not expressed (0) are represented with blue and grey squares, respectively. (D) Percentage of similarity between the model attractors under perturbations (vertical axis) and the binarized expressions of CLP, ETP, DN2A, DN2B, and DN3A cells (horizontal axis) (Jojic et al., 2013) (left). Description of known effect of the gene perturbation on T-cell development (right).

*Notch1*, a cell surface receptor that mediates Notch signaling, is known to play an essential role in early T-cell development. *Notch1* deficiency leads to blocked T-cell development and accumulation of other hematopoietic lineages (Radtke et al., 1999). Using our inferred executable network, we simulated the developmental trajectory of ETP cells in the absence of *Notch1*. Simulations predicted the presence of two possible steady state attractors, localized near the earlier section of the pseudo-time domain (**Figure 4B**). Comparing the expression states of the simulation attractors (**Figure 4C**) with the binarized expression of known cell states extracted from microarray data (Jojic et al., 2013), we found that the attractor states are more similar to ETP or CLP, and none show significant similarities to later stages of T-cell development (**Figure 4D**). This agrees with previous reports (Radtke et al., 1999) that lack of *Notch1* blocks T-cell development.

*Tcf7* is a crucial transcription factor for T-cell specification and differentiation that is upregulated by Notch signaling. Lack of *Tcf7* results in premature arrest of T-cell development before the DN2 stage (Weber et al., 2011). Our model predicts a single attractor state in the absence of *Tcf7* (**Figure 4B**). This attractor precedes the DN2 stage and does not express *Gata3, Bcl11b, Ets1, Cd3e*, or *Cd3g* (**Figure 4C**), in agreement with experimentally reported analysis of *Tcf7*-/-lymphoid-primed multipotent progenitors cultured *in vitro* (on OP9-DL4) at day 4 (Weber et al., 2011). The simulated *Tcf7* knockout steady state attractor also show more similarity to microarray profiles for ETP cells than either DN2 and DN3 cells (**Figure 4D**), which is in agreement with previous reports (Weber et al., 2011).

Next, we investigated the effect of simulated knock-out of *Bcl11b*, a crucial gene for T-cell commitment (Hosokawa et al., 2018). It has been shown experimentally that *Bcl11b* deficient cells cannot proceed beyond the DN2 stage (L. Li et al., 2010). Our *in silico* results predict one attractor in the absence of *Bcl11b* (**Figure 4B**). This attractor resembles the DN2A cell state (**Figure 4D**) which recapitulates the aforementioned experimental result of *Bcl11b* knockout (L. Li et al., 2010).

We also simulated the effects of perturbing *Runx1, Tcf12* and *Myb*. Knock-out of *Runx1* stops T-cell development before the DN3 stage (Egawa et al., 2007); knock-out of *Tcf12* results in developmental blockage before DP stage (Wojciechowski et al., 2007), whereas *Tcf12* overexpression enhances T-cell development(Braunstein & Anderson, 2012); knock-out of *Myb* causes multiple blocks during T-cell development (Bender et al., 2004). We have simulated these gene perturbations, and found qualitative agreement with the experimentally reported results (**Figure 4B, Figure 4C** and **Figure 4D**).

Next, we tested our model for a simultaneous perturbation of two genes. Interestingly, while the absence of Notch signaling results in loss of T-cell development, forced expression of *Tcf7* is able to partially rescue T-cell development in the absence of Notch (Weber et al., 2011). To test our model against this observation, we simulated shutting off the expression of *Notch1* (the mediator of Notch signaling) and forcing expression of *Tcf7* to the **‘**on**’** state simultaneously. This resulted in two attractors (**Figure 4B**), one of them localized at earlier stages, and one of them close to the DN2 stage, which reflects the limited but not complete development of T-cells when compared with the knock-out of *Notch1* (**Figure 4C** and **Figure 4D**).

In addition to these perturbations, we have performed full systematic GRN perturbations for one and two gene perturbations (**Figure S2A** and **Table S3** and **Table S4**) and sensitivity analysis of gene interactions (**Figure S2B, Table S3**, and **Table S4**). Taken together, we showed that our model can predict the effect of single and double gene perturbations on the developmental trajectory of early T-cell development.

## DISCUSSION

There is increasing availability of scRNA-seq datasets for different developmental systems. To date, the inference, analysis, and simulation of logical GRNs directly from scRNA-seq data have not been integrated together. Here we present IQCELL, an integrated strategy implemented as a Python package to infer, analyze, and simulate GRNs directly from scRNA-seq data and pseudo-time order of the cells.

IQCELL is able to capture, directly from scRNA-seq data, over 75% of the reported gene interactions in early T-cell development (Longabaugh et al., 2017). These interactions were obtained and characterized by decades of research and experiments (Yui & Rothenberg, 2014). For example, regulators of *Bcl11b*, an essential gene for T lineage commitment, were successfully identified by IQCELL. More than half of the interactions that have not been captured were *Spi1* effector genes, which is a Notch signaling antagonist (Rothenberg et al., 2019). However, Notch signaling contains *Spi1* inhibitory effect on T-cell regulators (Rothenberg et al., 2019) and potentially masks some of *Spi1* negative regulatory roles in early T-cell development.

We also tested the dynamics of the obtained GRN. We showed that when this logical GRN is simulated from ETP cell state, its dynamics evolves to the cell state associated with the DN3 stage, in agreement with experimental observations. Importantly, we showed that our platform can produce GRN models with high predictive power for the effect of genetic perturbations. For example, simulated knock-out of *Bcl11b* caused the developmental trajectory to halt at the DN2 stage, in agreement with experimental studies. We identified eight gene perturbations that halt T-cell development at different points between the ETP and DN3 stages, and IQCELL showed satisfactory agreement for all perturbations with experimental studies (**Figure 4C**).

These results show that the multi-step strategy implemented in IQCELL is effective for reconstructing functional GRNs from existing information in scRNA-seq data. Because its methodology is not specific to a single developmental system, IQCELL may be broadly useful in understanding how GRNs contribute to cell development in a variety of developmental contexts. IQCELL results may help uncover functional relations between genes and thereby help design more effective gene manipulation strategies to drive stem cell cultures toward fates of interest. Synthetic gene interactions can be added to the GRNs outputted by IQCELL to predict the effect of novel synthetic gene circuits on native cell GRNs. A major goal in systems biology is the creation of multi-scale models that connect the decisions of individual cells within a multicellular system to emergent properties of the whole tissue (Qu et al., 2011; Swat et al., 2012). IQCELL can fill an important layer in such multi-scale models. By exposing the intracellular decision-making machinery of single cells, IQCELL could interface with other methods that connects these cellular decisions to tissue-level dynamics.

To date, there have been many methods introduced for reconstructing GRNs from single-cell data (Pratapa et al., 2020) and many of them focus on finding some type of correlation between genes. In one case, binarized sc qPCR data was used to decode logical GRNs for embryonic blood development (Moignard et al., 2015). In another recent study, sc qPCR data and its pseudo-time order was used to decode GRNs of blood stem cells (Hamey et al., 2017). However, to the best of our knowledge, prior to IQCELL there has not been any existing method or platform to infer executable and logical GRNs from scRNA-seq data, nor have previous methods dealt with the associated challenges of lower sensitivity and dropout effects.

Although the IQCELL framework allowed us to effectively model regulatory modules as logical (Boolean) gates where no extra parameters are required, logical models are limited in some respects. Firstly, logical modeling cannot effectively capture dose-dependency in gene interactions; for example, it is known that the downstream responses to *Gata3* are dose-dependent (Rothenberg, 2019). We suggest in future that this aspect be captured by multilevel models (Collombet et al., 2017). Multilevel modeling requires more sensitive measurements of TF expression levels, which may become feasible with emerging TF profiling methods (Moffitt et al., 2016). Secondly, capturing biophysical timescales in the logical framework is not trivial; one solution would be assigning a weighted time scale (Sun et al., 2017) to each simulation update step of the logical model. This can potentially help to include some time-scale sensitive events in cell GRN dynamics, such as stochastic chromatin restructuring events (Ng et al., 2018).

Since IQCELL provides users with a flexible framework, future studies could integrate other sources of information such as binding of TFs to DNA via ChIP-seq (Johnson et al., 2007) and CRISPR screening on the effect of gene perturbations on developmental trajectories (Gilbert et al., 2014) to potentially improve GRN reconstruction. In a recent study, the combination of scATAC-seq and scRNA-seq with machine learning methods have been used to infer a set of informative transcription factors during differentiation (Kamimoto et al., 2020). In addition, new opportunities are arising to investigate the decision-making machinery of the cells in their native environment (via in-situ cell profiling) (Lee et al., 2015). The combination of these methods, prior knowledge of cell-cell interactions (Browaeys et al., 2019; Kirouac et al., 2009), and emerging theoretical knowledge and computational technologies for capturing and quantifying spatio-temporal information content of cell signaling (Cepeda-Humerez et al., 2019; Dubuis et al., 2013; Maity & Wollman, 2020; Ostblom et al., 2019) can be used as invaluable resources for the next generation of GRN inference methods. These next-generation methods would ideally integrate cell signaling (P. Li & Elowitz, 2019) with GRNs directly from multi-omics sc data. In conclusion, the results presented here suggest that IQCELL will be a broadly useful tool to study cellular decision making in a variety of developmental systems.

## AUTHORS CONTRIBUTIONS

Conceptualization, T.H. and P.W.Z; Methodology, T.H; Software, T.H, M.A.L, and A.Y.K Investigation, M.A.L, C.F, M.H, and S.S; Provision of study animals, K.M.M; Writing – Original Draft, T.H, D.A.H, and P.W.Z; Writing –Review & Editing,, T.H, D.A.H, M.A.L, K.M.M, C.F and P.W.Z; Supervision, P.W.Z;

## ACKNOWLEDGEMENTS

We thank Wen Zhou and Ellen V. Rothenberg for generating a publicly available high-quality scRNA-seq data set of early mouse T-cell development. We thank Sara-Jane Dunn, Boyan Yordanov, and Ellen V. Rothenberg for our fruitful discussions and Yale S. Michaels and John M. Edgar for critically reading the manuscript. We also thank Microsoft Research (Cambridge, UK) and Sara-Jane Dunn for facilitating the opportunity for the author to deepen his understanding of the Z3 reasoning engine. Funding: This work has been supported by CIHR Foundation and NSERC Discovery grants to P.W.Z.

## METHODS

### RESOURCES

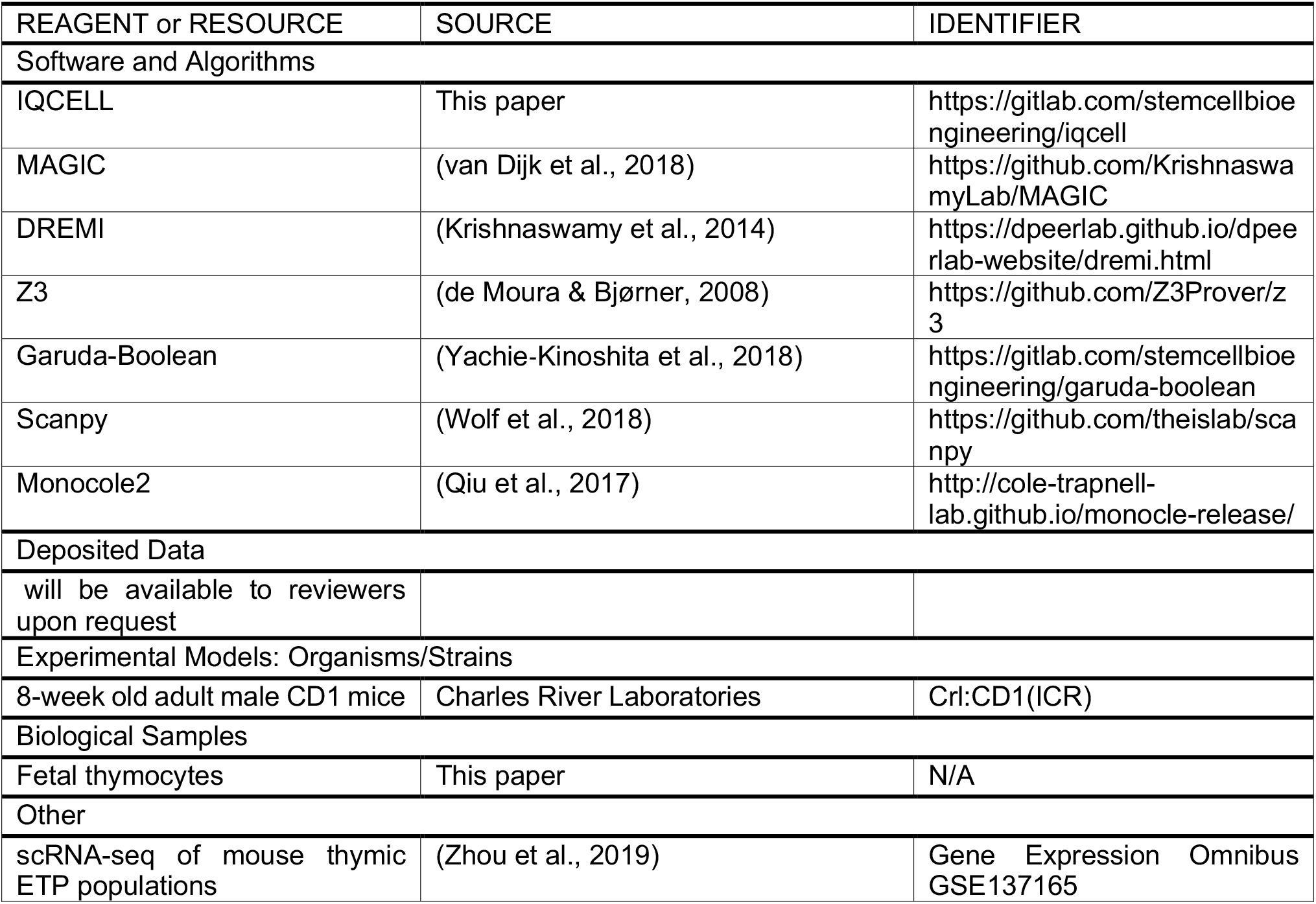

#### RESOURCE AVAILABILITY

##### Lead Contact

Further information and requests for resources and reagents should be directed to the Lead Contact, Peter Zandstra.

##### Data and Code Availability

The source code of IQCELL python package generated during this study is available on Gitlab: (https://gitlab.com/stemcellbioengineering/iqcell)

## METHOD DETAILS

### Gene expression recovery

scRNA-seq data is usually affected by dropouts, which is a technical term that is used to describe the false-negative reads of messenger RNAs. Dropout effects cause the expression profile of genes to be underrepresented. Usually, genes with low copy numbers (e.g transcription factors) are more affected by this effect. IQCELL applies a recent method (MAGIC) that uses a graph-based imputation method to infer the expression of dropouts (van Dijk et al., 2018) (**Figure S1A**). This imputation is important as dropouts can affect the inference of gene relations (**Figure S1A**).

### Gene selection

Selecting a small subset of genes to include in a functional GRN from the entire set of genes detected by scRNA-seq is, in general, a challenging task. Fortunately in mouse T-cell development, many relevant genes are known (Longabaugh et al., 2017). Here we describe a possible general approach for a selecting smaller subset of genes from a large set.

Selecting relevant genes for a functional/causal GRNs is generally a multilayered process (**Figure S1B**) which ideally combines multiple sources of information. One possible source is prior knowledge of important genes in the process, which can be obtained through literature review or systematic gene perturbation experiments (such as genome-wide CRISPR screening).. Alternatively, genes of interest can be selected directly from scRNA-seq data via various information theory metrics (Ang et al., 2016). Many scRNA-seq data analysis packages set an initial filter to only select highly variable genes (HVGs) for downstream analysis. HVGs are genes whose mean-scaled variance exceeds an automatic threshold. Although this is generally a useful filter, it does not necessarily select for genes whose expression levels vary significantly across pseudo-time. Therefore, IQCELL has a built-in function to visualize expression dynamics along pseudo-time and calculate the degree of variation, which can be used as an additional input for gene selection.

Beside these, there are other network-based approaches to select informative genes. These methods typically prioritize genes with connections to many other genes (high degree). Finally, enrichment analysis and ChIP-seq data can be another source of gene selection; however, these methods are generally low-throughput, noisy, and prone to false positives/negatives. For this study, we manually curated a list of genes based on biological significance (Longabaugh et al., 2017) and dynamics along the pseudo-time (**Figure S1B** and **Table S1**).

### Establishing the initial gene-gene interaction network

To form the initial gene-gene interactions network from scRNA-seq data (**Figure S1C**), IQCELL first forms a list of all possible pairwise gene-gene interactions. This list does not include autoregulation by default (optional). Next, it uses a recent method (DREMI) to calculate the resampled and conditional mutual information between gene pairs (Krishnaswamy et al., 2014) (**Figure S1C**). In general, the mutual information (I) between a pair of variables X and Y (where X and Y represent two genes) is calculated as below:

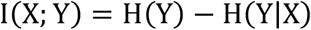

where H(X) is called the entropy of distribution X and P(X) is the probability density of the variable X:

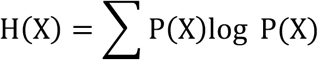

The mutual information score is symmetric and unsigned. Next, IQCELL applies the Pearson correlation coefficient to assign a sign (+/-which represents activation/repression) to the interactions, based on the sign of the correlation between two genes (**Figure S1C**):

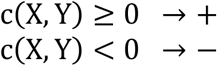

Where c(X,Y) is the Pearson correlation coefficient of the pair of gene X and Y:

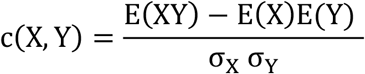

E(X) is the expected value and σ_X_ is the standard deviation of X. Finally, IQCELL eliminates interactions with mutual information values smaller than a pre-defined threshold (with the default value of one standard deviation below the mean). The result is an undirected and signed interaction network.

### Binarization of gene expressions

IQCELL binarizes the expression for each gene individually. To do this, it can apply two possible methods that can be chosen by the user (**Figure S1D**). The first method finds the mean of expression and assigns **‘**off**’** (0) to the genes (for each cells) with the expressed numbers of mRNAs that are smaller than the prefixed threshold and **‘**on**’** (1) if they are larger. The second method (the default method) binarizes the expression based on the cut-off identified by k-means clustering (k=2) method (**Figure S1D**)(Macqueen, 1967). In general, we group the expression of each gene in each cell g_i_ (i ∈ N_cells_) into two possible groups S_i_ = {S_off_, S_on_} by finding:

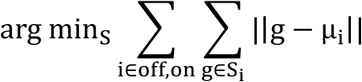

In which µ_i_ (centroids) is the mean of expression values in **‘**on**’** or **‘**off**’** cluster. The threshold (τ) is obtained as the average of two centroids:

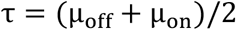

The binarized gene expression value (G_i_) is obtained similar to the mean method:

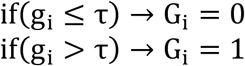

### Establishing the gene hierarchy via pseudo-time

After binarization of expression levels, IQCELL calculates an interaction hierarchy to further filter gene-gene interactions, thereby eliminating interactions that are less likely to be causal. First, it averages the binarized expression values in a sliding window over pseudo-time (**Figure S1E**). To do this, IQCELL first sorts the cells based on their pseudo-time values **c**_i_ (i ∈ N_cells_). Next, it averages the values of binarized gene expressions over an averaging window (with the default length of L = N_cells_/N_genes_). This results in a density representation of the binarized gene expression values along pseudo-time (t) (**Figure S1E**):

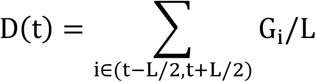

Next, it calculates transition points between low-to-high or high-to-low density regions for all genes (**Figure S1E**) and sorts genes based on their transition points (**Figure S1E**). Finally, IQCELL includes autoregulation (autoactivation) as a possible self-interaction for genes with less than two possible activators. This step is optional and is used in the current study, leading to inclusion of autoactivation interactions for *Hes1, Notch1, Lmo2*, and *Spi1* in the final interaction network.

### Implementing the Z3 reasoning engine to infer logical GRNs

To generate functional GRNs, IQCELL implements a modified network inference strategy in the Z3 engine (Hamey et al., 2017). This method effectively finds optional logical rules for each gene based on the possible list of interactions obtained from previous steps. The optimal rules are those that when executed as logical gates for each gene (given the state updates of other genes along the pseudo-time as an input), follow the experimental data. This is quantified with the percentage of similarity (based on Hamming distance) between the two (**Figure S1F**). Similar to (Hamey et al., 2017), we allow up to four possible activators and up to two repressors for each gene. In contrast to (Hamey et al., 2017) and similar to (S.-J. Dunn et al., 2014), for gene activation, we assume that all the activators are necessary (which is implemented with the **‘**and**’** logic gate), but only one repressor is enough for repression (which is implemented with the **‘**or**’** logic gate). In summary, the most general logical rule for the regulation of a gene (g_j_) by (the maximum number of) six regulators including (maximum) four activators (A_1_, A_2_, A_3_, A_4_) and (maximum) two repressors (R_8_, R_4_) is:

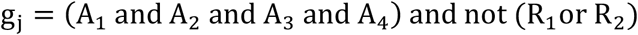

Which indicates that all activators and none of the repressor should be expressed for the gene to be expressed. Next, for each gene, the rule with the highest average mutual information for interactions is selected by IQCELL for the final GRN (**Table S1**).

### Asynchronous simulator of GRN under normal and perturbed condition

To analyze the system-level behavior of the obtained GRN models and predict the effect of gene perturbations on developmental trajectories, IQCELL uses (asynchronous) Boolean simulations. Boolean GRNs contain a set of genes 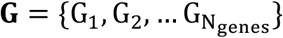 and their update function which encodes the gene regulatory details (one update function per gene). The update function of a gene 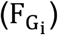 implies what will be the activity state of that gene (on or off) at the next discrete time point G_i_(t + 1) given the state of all the genes at the current discrete time point 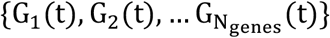:

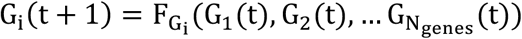

IQCELL, uses the asynchronous update strategy (Yachie−Kinoshita et al., 2018). In this strategy, at each discrete time step, only one random gene is selected and updated. This results in stochastic dynamics. This lets us average the expression states (average_exp) at each discrete time point (j) over the ensemble of stochastic states that started from the same initial point and are now at that particular time point (**EX**(j)) (**Figure 3G**):

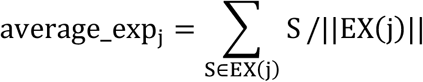

Where S = {**0**,1} therefore:

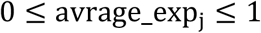

IQCELL also has a built-in function to perturb the GRNs in two ways. It has the capability to perform KO (setting the gene to be always **‘**off**’**) and OE (setting the gene to be always **‘**on**’**) experiments simultaneously for one or multiple genes systematically. Also it can perturb the gene-gene interactions systematically (**Figure S2**).

### Software architecture of IQCELL

IQCELL is implemented as a Python package. It is modular and scalable, to help researchers to expand, optimize, and customize it for their future studies. Also, it has a minimal Python interface, which allows the users to use it with minimal computational skills, and implement it to their system of interest.

### Pre-processing the scRNA-seq data

To dissect the developmental trajectory of T-cell development (from ETP to DN3 stages), we used a scRNA-seq dataset from the Rothenberg Lab (Zhou et al., 2019). We used the analysis pipeline provided by the Theis Lab (Luecken & Theis, 2019). After the quality control and normalizing expression values, single-cell transcriptional states were visualized in reduced dimensional space using UMAP (**Figure S4A**). To understand the underlying structure of data we perform clustering based on the Louvain method (**Figure S4B**) which yields 14 sub-clusters. Next, we evaluate the expression pattern of developmentally important genes in blood and particularly T-cell development. ETP associated genes (*Flt3, Lmo2, Mef2c*) are all expressed in the cell clusters in the left side (**Figure S4C**). DN2-a stage is marked by *Il2ra*, this gene along with the gene associated with committed DN2 cells (*Bcl11b*) and DN3 associated genes (*Cd3e* and *Rag1*) expressions are localized on the right side (**Figure S4C**). Granulocyte lineage marker (*Elane*) and macrophage lineage marker (*Mpo*) expression are high at cluster 13 (not shown) and we excluded this cluster for future analysis. This can be due to alternative lineage decisions in development or contamination. Altogether we conclude that cluster 0 includes many of the cells at the ETP stage and the developmental progression is toward clusters 9 and 11 as the endpoints (**Figure S4C**).

As previously reported (Zhou et al., 2019), we use a supervised approach in pseudo-time ordering on the subset of genes that are known to be developmentally important in T-cell development or are alternate lineage markers. The selected genes were similar to (Zhou et al., 2019) and DDRtree pseudo-time ordering is performed on the data (Qiu et al., 2017). As established in the cluster analysis, step cluster number 0 is the best candidate as the cluster with the earliest developmental stage and is used as the root for the algorithm. The result shows a single trajectory starting at cluster 0 and progressing toward later stages (**Figure S4D**). Pseudo-time ordering shows the dynamics of genes as a function of differentiation (**Figure S4E**).

### Relationship between the quality of single cell data and the GRN inference

As a short note, there is a direct relationship between the quality of the scRNA-seq data and inferred pseudo-time with the quality of the inferred GRN. A suitable scRNA-seq data has high resolution with regard to the underlying expression dynamics during the developmental process of interest, sufficient read depth, and yields a pseudo-time trajectory that qualitatively resembles the known gene expression state progression of the developmental system to be modeled (Zhou et al., 2019).

### Sample preparation and Single-cell RNA-sequencing of in vitro T-cell differentiation

Isolated fetal liver cells from decapitated E13.5 CD1 mouse embryos were subjected to TER-119 depletion by EasySep magnetic sorting (STEMCELL Technologies). Next, sorted HSPCs (Sca-1+ cKit+) cultured at 3.1 x103 HSPCs/cm2 (corresponding to 1000 cells/well) in DL4 (10 μg/mL) and VCAM-1 (2.32 μg/mL) coated 96-well plates (Shukla et al., 2017).10X Chromium was used to prepare single-cell cDNA libraries, and Illumina Nextseq was used to 3**’** sequence the samples. Gene-barcode expression matrices were calculated from the raw data via CellRanger (10X Genomics).

## SUPPLEMENTAL FIGURES

**Supplementary Figure S1.**
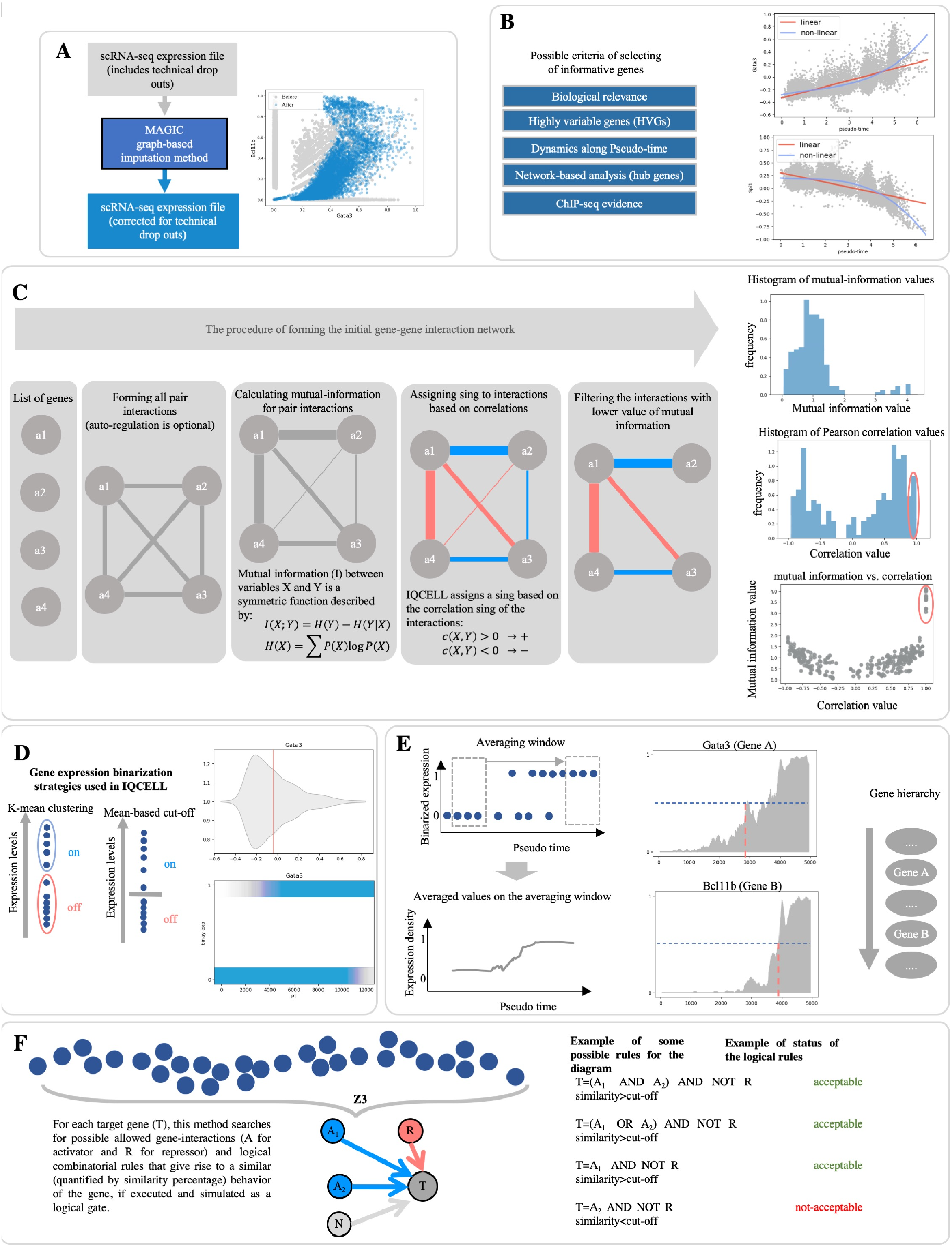
Overview of IQCELL parts and algorithms (related to Fig. 1). (A) Overview of gene expression recovery step. The scRNA-seq data is corrected for dropout effect via a borrowed library **‘**MAGIC**’** from literature (left). Raw (grey) and recovered (blue) expression of Bcl11b vs. Gata3 (right). (B) Overview of gene selection step. The most important criteria for gene selection is biological relevance and literature curation. However, the user can benefit from observing the dynamics along the variability pseudo-time and variability of genes. Other network-based methods and chip-seq evidence can be used as supplementary methods for gene selection (left). To visualize the dynamics of genes along the pseudo-time (right), IQCELL has a built-in function to visualize and fit linear and non-linear functions to the gene. vs time function. Users can use this information to select genes as well. (C) Overview of generating the initial interaction network. The steps toward obtaining the directional and signed interaction network from an initial list of genes (left column). The histogram of mutual information between gene pairs, the histogram of Pearson correlation between gene pairs (the correlation value of 1 is for the correlation of genes with themselves, marked by red), and mutual information vs. Correlation values (right column). (D) Overview of expression binarization step. There are two implemented binarization methods in IQCELL. K-means clustering (default) and binarization based on the mean value of expression of the gene between all the cells (left). Example of binarization of genes and their expression along the pseudo time (right). (E) Overview of generating the gene hierarchy step from the binarized gene expressions. First, the expression levels are averaged with a sliding window along the pseudo-time. This results in the density profile of binarized genes along the pseudo-time (left). Next, based on clustering the density, the transition point (from high to low or low to high) are captured (center). Finally, genes are sorted based on transition points. Genes can interact with genes with a lower rank (right). (F)Overview of implementing the Z3 reasoning engine. At this step, the filtered set of interactions are used to make provisional update rules for each gene.

**Supplementary Figure S2.**
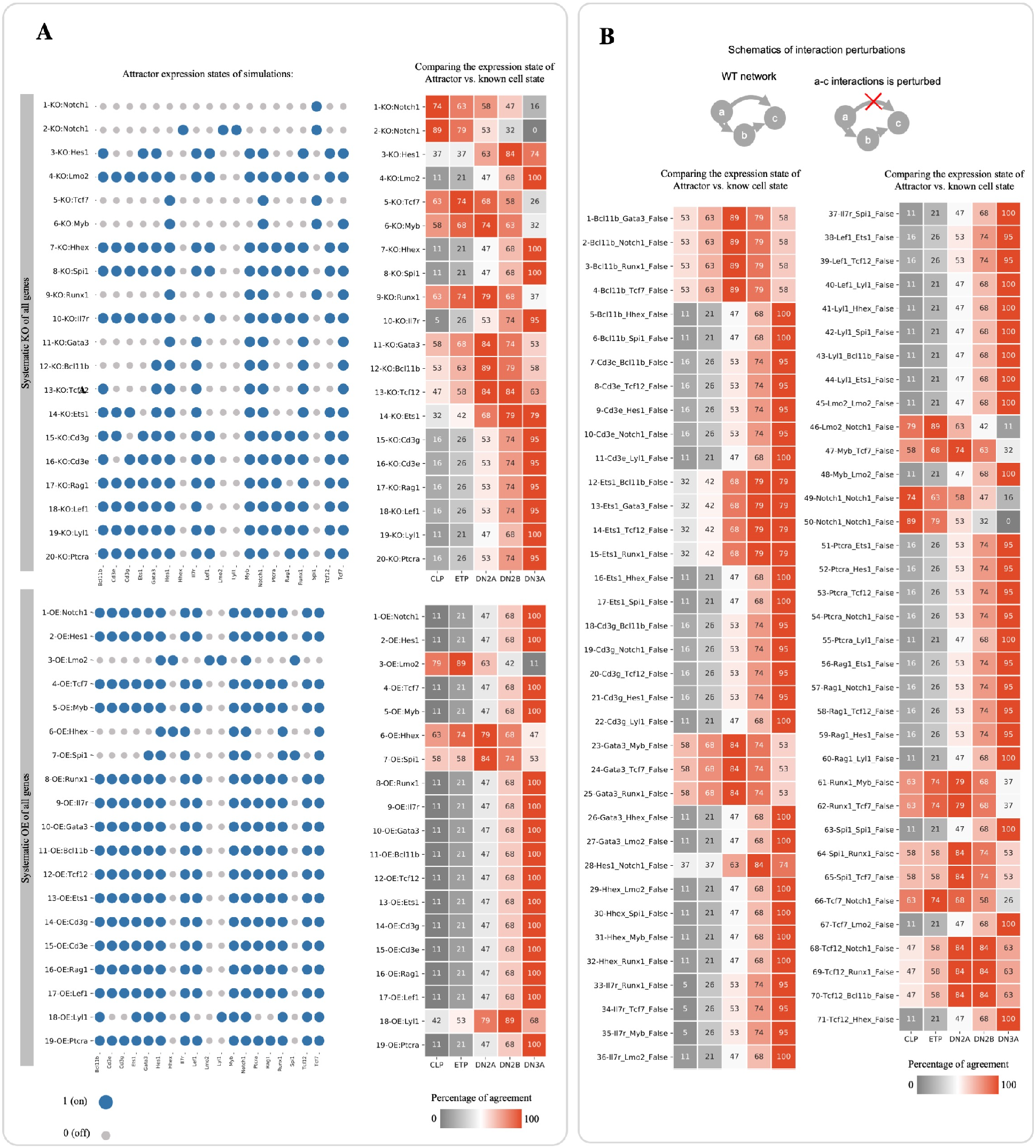
GRN perturbations via IQCELL(related to Fig. 4). (A) Systematic gene perturbations. The expression states of the model attractor (left). The percentage of similarity between the attractors (vertical axis) and the binarized expressions of CLP, ETP, DN2A, DN2B, and DN3A cells (horizontal axis) (Jojic et al., 2013) (right) for systematically perturbed GRN with single gene KO and OE. (B) Systematic gene-gene interaction perturbations. Overview of GRN link perturbation (top). The percentage of similarity between the attractors (vertical axis) and the binarized expressions of CLP, ETP, DN2A, DN2B, and DN3A cells (horizontal axis) (Jojic et al., 2013) (bottom) for systematically perturbed GRNs.

**Supplementary Figure S3.**
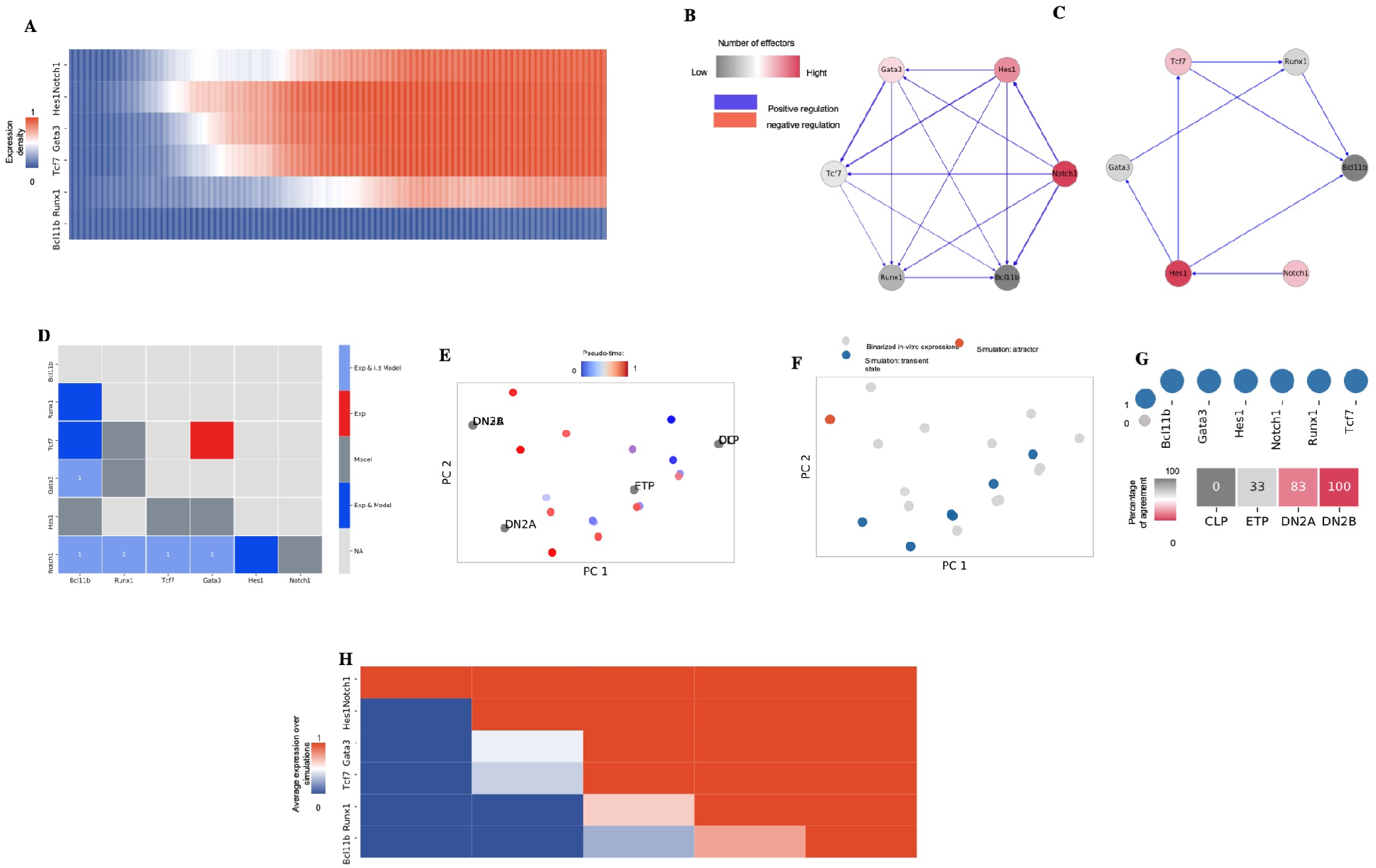
Implementation of IQCELL with another early T-cell development scRNA-seq data set. (related to Fig. 3) (A) To demonstrate the universality of IQCELL, we have tested this planform with another in-house scRNA-seq data. The IQCELL tutorial in the IQCELL website is based on this dataset. We used 10X scRNAseq by performing whole-genome transcriptional analysis. These experiments are performed with the mouse T-cell progenitor populations from fetal liver (FL) hematopoietic stem and progenitor cells (HSPCs) differentiated in vitro using the DL4+VCAM platform(Shukla et al., 2017). FL HSPCs were seeded on DL4+VCAM coated plates and cultured for 4 or 7 days prior to analysis, or immediately sorted and captured for library preparation. Pooling cells from multiple differentiation time points enabled sampling of cells from the entire T-cell lineage progression, rather than just endpoint transcriptional states. Here we have selected a small set of 6 genes that are important in the early T-cell development (from ETP to DN2 stages). The heat map shows the expression matrix of smoothed binarized expressions along the pseudo-time. (B) The set of all possible gene-gene interactions, filtered by interaction hierarchy and mutual information cut-off (the thickness of lines represents the mutual information between genes), and signed by correlation. (C) Provisional GRN for early mouse T-cell development obtained by Z3 step with additional maximize mutual information criteria. (D) Detailed representation of the proposed interactions provided by IQCELL and experimentally reported ones. (E) PCA plot of the binarized scRNA-seq data color-coded with the pseudo-time values attributed to each cell. The binarization is performed by clustering the cs RNA-seq expressions into expressed or not expressed levels. On top of that, the binarized expressions of CLP, ETP, DN2A, DN2B, cells have been calculated from the immgen microarray data (Jojic et al., 2013) and overlaid on RNA-seq data. (F) PCA plot of the simulated developmental trajectories are overlaid on the binarized scRNA-seq. (G) Expression states of the model attractors (top). The percentage of similarity between attractor and known cell states (Jojic et al., 2013). (H) Averaged gene expression of the simulated data at each simulation step. All simulations started from the same initial condition.

**Supplementary Figure S4.**
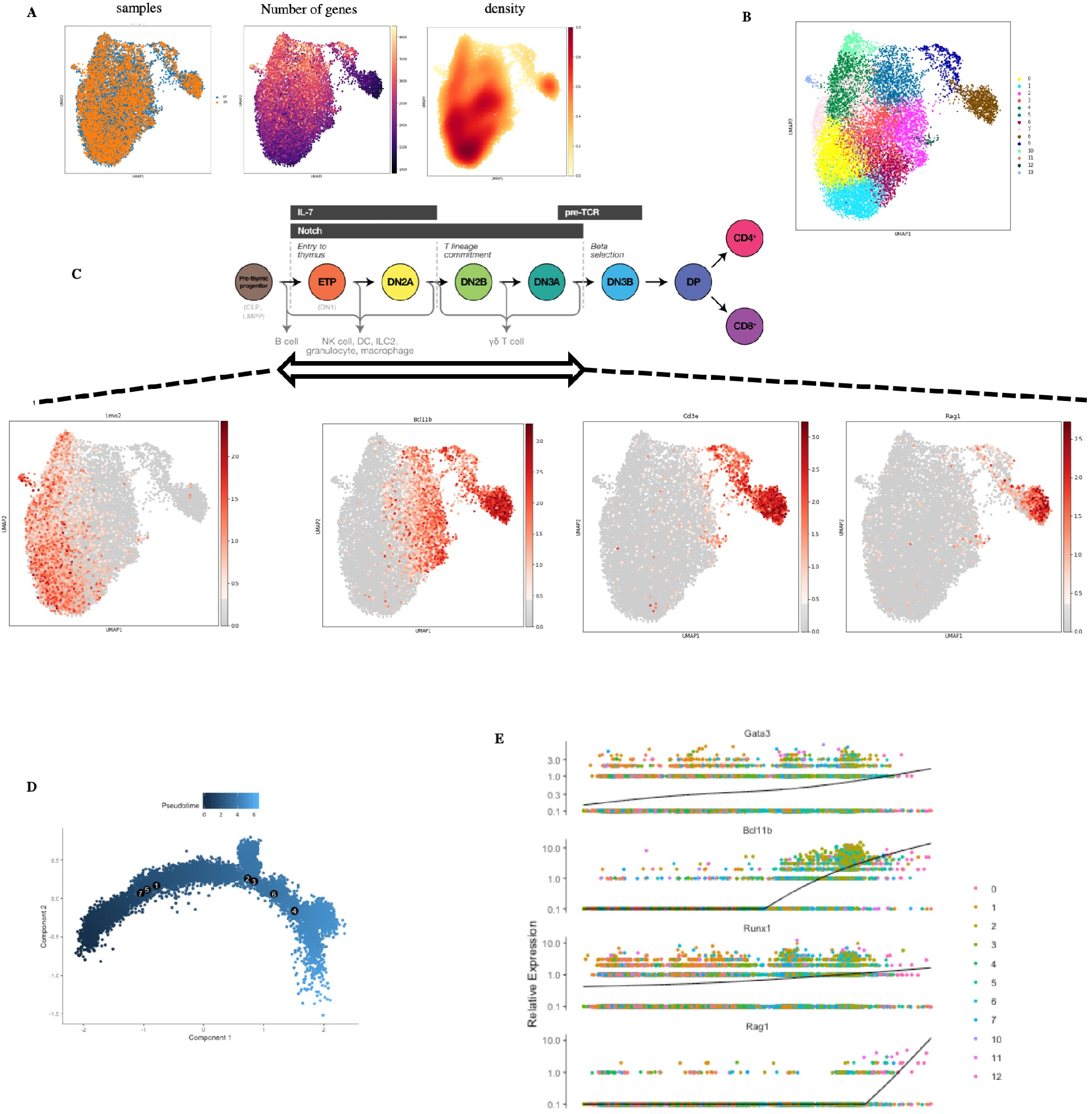
The overall workflow of preprocessing the scRNA-seq data (related to Fig. 3). **(A)** UMAP representation of Data overlaid with sample id, number of genes per cell, and density plot. (B) Clustering of scRNA-seq data. (C) Gene expression of stage-specific gene overlaid on top of UMAP. (D) The pseudo-time trajectory of the data inferred by Monocle platform. (E) Example expression of genes along the pseudo-time trajectory.

## SUPPLEMENTAL TABLES

**Supplementary Table S1.**
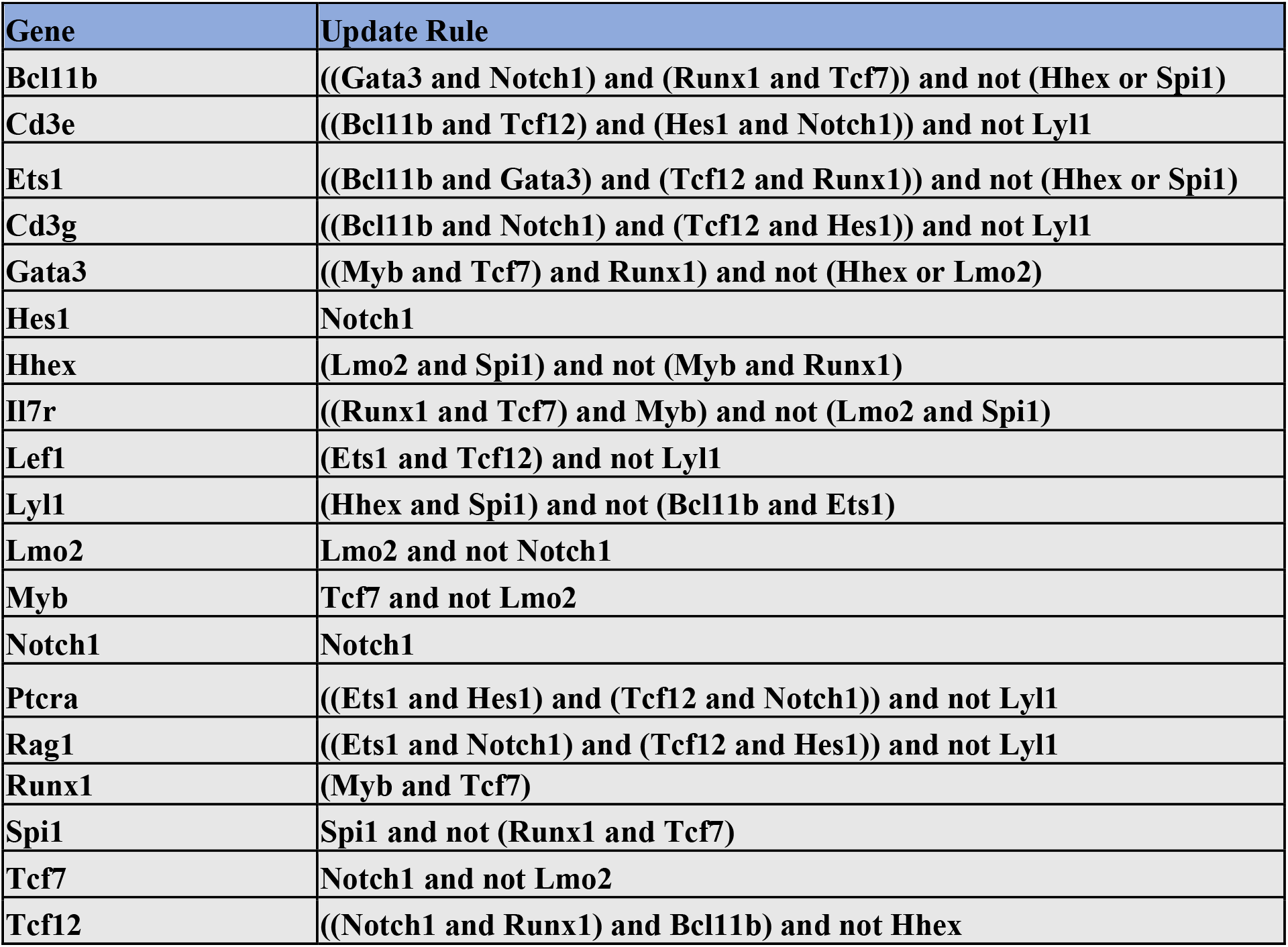
(related to Fig. 2 and 3). The final gene list and provisional logical GRN for early T-cell development. The rules are picked from possible rules in Z3 step based on maximizing the average mutual information per gene.

**Supplementary Table S2.** (related to Fig. 2). Genes used for supervised trajectory inference.

| Gene |  |
| --- | --- |
| Bcl11a | Itgax |
| Bcl11b | Kit |
| Ccr9 | Lef1 |
| Cd34 | Lmo2 |
| Cd3e | Ly6d |
| Cd3g | Lyl1 |
| Cd44 | Mef2c |
| Cd82 | Meis1 |
| Cebpa | Mpo |
| Cxcr4 | Myc |
| Dtx1 | Mycn |
| Erg | Nfil3 |
| Ets1 | Notch1 |
| Ets2 | Nrarp |
| Flt3 | Nt5e |
| Gata1 | Pdgfrb |
| Gata2 | Pgk1 |
| Gata3 | Pim1 |
| Gfi1 | Ptcra |
| Gfi1b | Rag1 |
| Hes1 | Rag2 |
| Hhex | Runx1 |
| Hoxa9 | Runx2 |
| Id2 | Runx3 |
| Id3 | Sox13 |
| Ikzf1 | Spi1 |
| Ikzf2 | Spib |
| Il2ra | Tcf12 |
| Il4ra | Tcf7 |
| Il7r | Tlr7 |
| Irf8 | Zbtb16 |
| Itga2b | Zfpm1 |
| Itgam |  |

Supplementary Table S3. (related to Fig. 4). Comparison of attractors of perturbed GRN with the known cell states from microarray data. (Online)

Supplementary Table S4. (related to Fig. 4). The attractor states of perturbed GRN. (Online)

